# Beyond accuracy: Measures for assessing machine learning models, pitfalls and guidelines

**DOI:** 10.1101/743138

**Authors:** Richard Dinga, Brenda W.J.H. Penninx, Dick J. Veltman, Lianne Schmaal, Andre F. Marquand

## Abstract

Pattern recognition predictive models have become an important tool for analysis of neuroimaging data and answering important questions from clinical and cognitive neuroscience. Regardless of the application, the most commonly used method to quantify model performance is to calculate prediction accuracy, i.e. the proportion of correctly classified samples. While simple and intuitive, other performance measures are often more appropriate with respect to many common goals of neuroimaging pattern recognition studies. In this paper, we will review alternative performance measures and focus on their interpretation and practical aspects of model evaluation. Specifically, we will focus on 4 families of performance measures: 1) categorical performance measures such as accuracy, 2) rank based performance measures such as the area under the curve, 3) probabilistic performance measures based on quadratic error such as Brier score, and 4) probabilistic performance measures based on information criteria such as logarithmic score. We will examine their statistical properties in various settings using simulated data and real neuroimaging data derived from public datasets. Results showed that accuracy had the worst performance with respect to statistical power, detecting model improvement, selecting informative features and reliability of results. Therefore in most cases, it should not be used to make statistical inference about model performance. Accuracy should also be avoided for evaluating utility of clinical models, because it does not take into account clinically relevant information, such as relative cost of false-positive and false-negative misclassification or calibration of probabilistic predictions. We recommend alternative evaluation criteria with respect to the goals of a specific machine learning model.

## Introduction

Machine learning predictive models have become an integral method for many areas of clinical and cognitive neuroscience, including classification of patients with brain disorders from healthy controls, treatment response prediction or, in a cognitive neuroscience setting, identifying brain areas containing information about experimental conditions. They allow making potentially clinically important predictions and testing hypotheses about brain function that would not be possible using traditional mass univariate methods (i.e., effects distributed across multiple variables). Regardless of the application, it is important to evaluate the quality of predictions on new, previously unseen data.

A common method to estimate the quality of model predictions is to use cross-validation and calculate the average prediction performance across test samples (Varoquaux et al., 2017) or balanced variants that account for different class frequencies (Brodersen et al., 2010) Selection of appropriate performance measures is a widely studied topic in other areas of science, such as weather forecasting (Mason, 2008), medicine (Steyerberg et al., 2010) or finance ((Hamerle et al., 2003). However, in the neuroimaging field, this has received surprisingly little attention. For example, many introductory review and tutorial articles focused on machine learning in neuroimaging (Haynes and Rees, 2006; Pereira et al., 2009; Varoquaux et al., 2017) do not discuss performance measures other than accuracy (i.e., simple proportion of correctly classified samples). Accuracy is also the most frequently reported performance measure in neuroimaging studies employing machine learning, even though in many cases alternative performance measures may be more appropriate for the specific problem at hand. However, no performance measure is perfect and suitable for all situations and different performance measures capture different aspects of model predictions. Thus, a thoughtful choice needs to be made in order to evaluate the model predictions based on what is important in each specific situation. Different performance measures also have different statistical properties which need to be taken into account. For example, how reliable or reproducible are the results, or what is the power of detecting a statistically significant effect?

In this paper, we provide a didactic overview of various performance measures focusing on their interpretation and practical aspects of their usage. We divide the measures into four families based on what aspects of a model prediction they evaluate; (1) measures evaluating categorical predictions, (2) ranks of predictions, (3) probabilistic predictions with respect to a squared error, and (4) probabilistic predictions with respect to information criteria. This includes accuracy, the area under the receiver operating characteristic curve, the Brier score, and the logarithmic score respectively, as most prominent members of each family. Next, we perform an extensive empirical evaluation of statistical properties of these measures, focusing on power to detect a statistically significant effect, power to detect a model improvement, evaluation of stability of the feature selection process, and evaluation of the reliability of results. We show that accuracy performs the worst with respect to all examined statistical properties. Last, we discuss appropriate evaluation criteria with respect to various goals of the specific machine learning model.

### Tutorial on performance measures

This section is a didactic overview of 4 main types of performance measures.

#### Categorical measures

The most commonly used performance measures are based on an evaluation of categorical predictions. These can be derived from a confusion matrix where predicted labels are displayed in rows and observed labels are displayed in columns (Figure 1). The most commonly used measure is accuracy, which is a simple proportion of all samples classified correctly. This can be misleading in imbalanced classes, since for example if the disease prevalence is 1% it is trivial to obtain an accuracy of 99% just by always reporting no disease. For this reason, balanced accuracy is often used instead (Brodersen et al., 2010). Balanced accuracy is simply the arithmetic mean of sensitivity and specificity, thus it is equal to accuracy when the class frequencies are the same (Figure 1), otherwise, because sensitivity and specificity are contributing equally, it weighs false positive and false negative miss classifications according to class frequencies.

**Figure 1:**
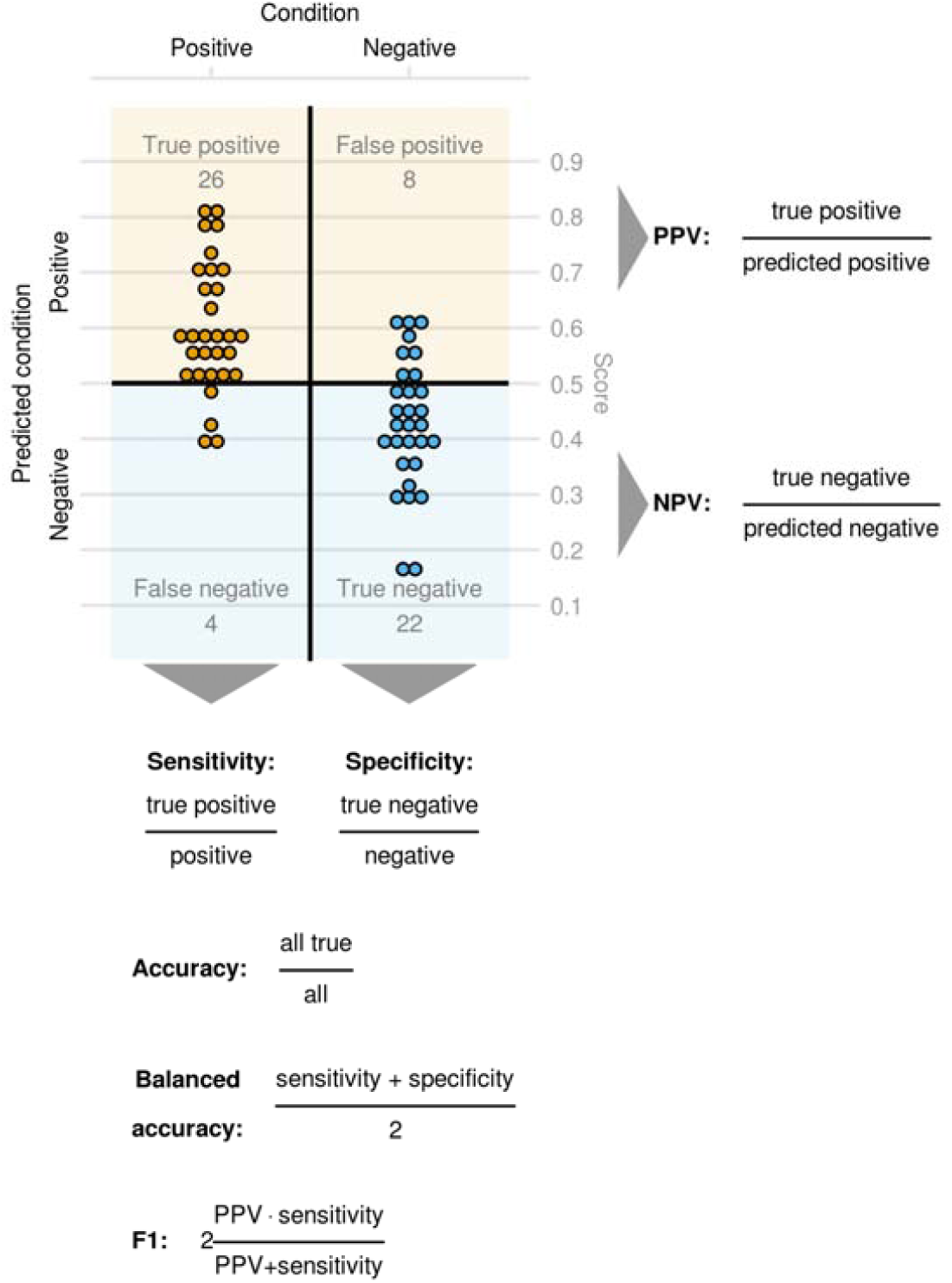
Definition of categorical predictions and their relationship to a confusion matrix. Note that the predicted score (e.g. predicted probability from a logistic regression, here depicted by the position of the predictions on the y-axis) needs to be dichotomized into categorical predictions, thus the magnitude of a miss-classification is not taken into account. Balanced accuracy equals accuracy when the class frequencies are equal, otherwise misclassification of the minority class is weighted higher, therefore, the chance level stays at 0.5. PPV: positive predictive value. NPV: Negative predictive value.

Sensitivity (true positive rate, recall) is the proportion of positive samples classified as positive, or in other words, it is the probability that a sample from a positive class will be classified as positive. Specificity (true negative rate) is the counterpart to sensitivity measuring the proportion of negative samples correctly classified as negative. Sensitivity and specificity do not take class frequencies (and class imbalances) or disease prevalence into account and they do not capture what is important in many practical applications. For example, in a clinical setting, we don’t know if the patient has a disease or not (otherwise, there would be no need for testing).Instead, what is known are the results of the test (positive or negative) and we would like to know, given the result of the test, what is the probability that the patient will have a disease or not. This is measured by positive predictive value (PPV) and negative predictive value (NPV) for positive and negative test results, respectively. It is easy to misinterpret sensitivity as PPV, the difference is subtle but crucial. To illustrate, say we have the following confusion matrix:

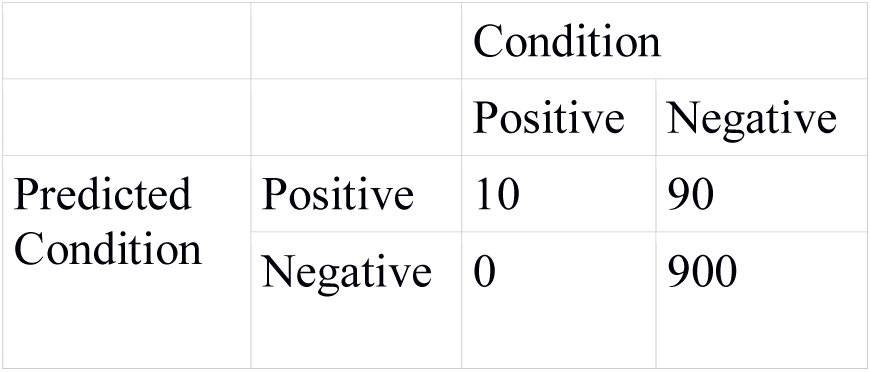

This gives us 100% sensitivity (10/10, 91% specificity (900/990), 91% accuracy ((10+900)/1000) and 96% balanced accuracy ((10/10 + 900/990)/2). According to these measures, the model performs well. However, in this example the disease prevalence is 1% (10/1000), so only 10% of patients with a positive test result are truly positive, the remaining 90% are misclassified (i.e., cell with positive predicted condition but negative actual condition in confusion matrix). Therefore, the test may actually be useless in practice. Another measure used in cases with imbalanced classes is the F1 score, which is the harmonic mean of PPV and sensitivity (so if one of the elements is close to 0, the whole score will be close to 0) (Figure 1). Since the harmonic and not the arithmetic mean is used, both sensitivity and PPV need to be high in order for the score to be high. Thus, this score emphasizes the balance between sensitivity and PPV and it assumes that both are equally important.

All categorical measures suffer from two main problems: first, they depend on an arbitrarily selected classification threshold. Each data point counts either as a correct or incorrect classification, without taking the magnitude of the error into account. If the decision threshold is set to 50%, then if a model predicts that a disease probability in a healthy subject is 49% percent and thus classifies this subject as healthy, this is indistinguishable from a disease probability of 1% in another healthy subject (also classified as healthy). Imagine two models, one predicts that a healthy subjects has a disease with a probability of 99% and other with a probability of 51%. The latter model is obviously better since the error is smaller, but if these predictions are thresholded at traditional 50% percent, the accuracy will not be able to detect the improvement. Due to this insensitivity, compared to measures that do not require dichotomization of predictions, using accuracy necessarily leads to a significant loss of statistical power. Second, false positive and false negative misclassification are weighted as being equally bad or according to class frequencies (balanced accuracy), which is often inappropriate. Rather, misclassification costs are asymmetric and depend on consequences of such misclassification. For example, in a clinical context, misclassifying a healthy subject as diseased is worse when this misclassification will lead to an unnecessary open brain surgery, than when it will lead to a prescription of medications with no side effects.

One way to combat unequal misclassification cost is to select the decision threshold according to cost-benefit analysis, such that the action is made when the expected benefit of the action outweighs the expected harm. This is then evaluated by cost-weighted accuracy, where true positives and false positives are weighted according to their relative cost. Although this allows to evaluate categorical predictions according to their utility and not arbitrarily, the dichotomization still creates problems for many practical applications. For example, in a clinical setting, the categorized predictions hide potentially useful clinical information from decision-makers (be it clinician, patient, an insurance company, etc.). If a prediction is only the presence versus the absence of a disease, a decision maker cannot take into account if a probability of the disease is 1%, 15% or 40%. This effectively moves the decision from stakeholders to a data analyst assuming that the chosen decision threshold is appropriate and constant for every situation and every patient, thus not allowing to take clinicians’ opinions or patients’ preferences into account.

#### Rank based measures

Another important family of performance measures are those based on ranks of predictions. Here, predictions are not categorical, but all predictions are ranked from lowest to highest with respect to the probability of an outcome.

The most common measure from this family is the area under the receiver operating characteristic curve (AUC). It measures a separation between two distributions, with a maximum score of 1 meaning perfect separation or no overlap between distributions and 0.5 being chance level when the distributions cannot be separated. In the case of model evaluation, the two distributions are distributions of predicted values (e.g. probability of a disease) for each target group. AUC is identical or closely related to multiple more or less known concepts, some of which we will review below.

The most common way of interpreting the AUC is through the receiver operating characteristic curve (ROC) (Figure 2A). This is a plot showing how a proportion of false positive and false negative misclassifications (i.e. sensitivity and specificity) changes as a function of the decision threshold. AUC is then the area under this curve. The curve itself is useful because it visualizes sensitivity and specificity across all thresholds not only for one threshold as in the case with the confusion matrix. Thus it allows choosing a decision threshold with an appropriate balance between sensitivity and specificity. We can see that if a model has no predictive power, then regardless of the threshold, the proportion of false positives and false negatives will always sum to 1, therefore the ROC curve will be a straight line across the diagonal and thus the area under this curve will be 0.5.

**Figure 2.**
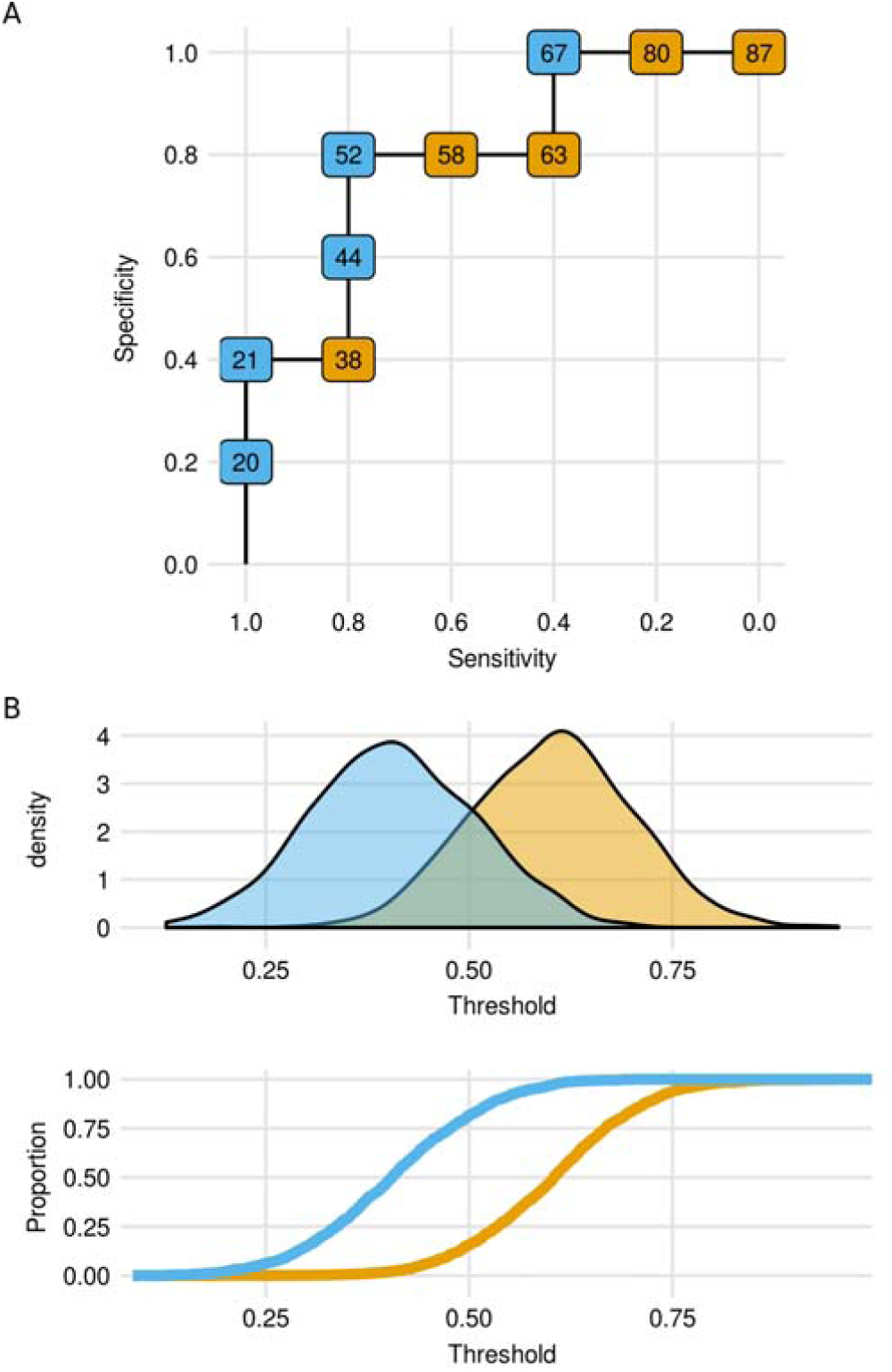
A: ROC curve for the same data as in Figure 3. The points on the curve represent threshold values that divide continuous prediction into two classes (represented as orange and blue colors). B: AUC as an overlap between two distributions. Two distributions (top) are transformed into cumulative distribution (bottom). The non-overlapping area of these two distributions is the area between two cumulative distribution curves, which equals to the area between the diagonal to the ROC curve or AUC – 0.5.

AUC can also be seen as quantifying to which extent two distributions overlap. We can rearrange the ROC plot, in such a way that the threshold value is on the x-axis and two curves are shown, one for the false-negative rate (1-sensitivity) and one for the true negative rate, both on the y-axis. The area between the diagonal and the ROC curve in the ROC plot (Figure 2A) is now the area between these two curves. These two curves are cumulative distributions of subjects from each class. The area between these curves represents the non-overlapping areas of two distributions (Figure 2B).

AUC is identical to C-index or a concordance probability in the case of a binary outcome (Hanley and McNeil, 1982). This is the probability that a randomly chosen data point forms a positive class is ranked higher than a randomly chosen example forms the negative class. E.g. if we have two patients, one with disease and one without, AUC is the probability that the model will correctly rank patients with a disease to have a higher risk of the disease than patients without the disease. This can also be interpreted as a proportion of all pairs of subjects in the dataset, where a subject with the disease is ranked higher than a subject without the disease (Figure 3B).

**Figure 3.**
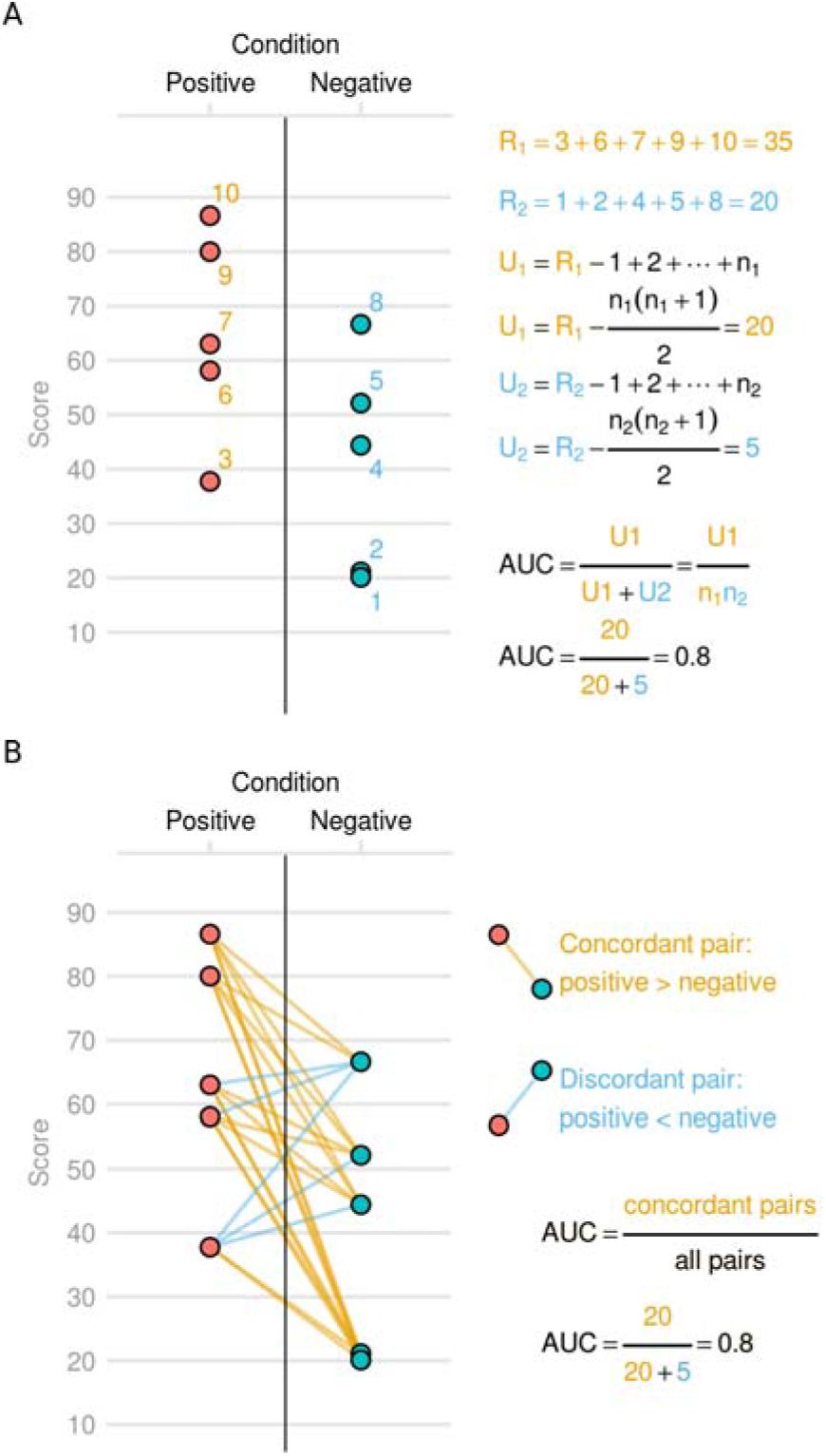
Interpretation and construction of AUC from the figure 2A as an **A) sum of ranks, i.e. Mann-Whitney U statistics.** Predictions are ranked from lowest to highest, and the sum of ranks R1 for the positive class is computed (or R2 for negative class). From the sum of ranks, Mann-Whitney U statistic is computed by subtracting the sum of ranks of one group from the sum of ranks of all predictions. AUC is the normalized version of the Mann-Whitney U, by dividing U statistics by the maximum possible value of the U statistics and **B) concordance measure.** AUC can be interpreted as a proportion of pairs of subjects where a subject from the positive class is ranked higher than a subject from the negative class, or where a randomly selected subject from the positive class is ranked higher than a randomly selected subject from the negative class.

AUC is also a rescaling of several rank-based correlation measures, including Sommers d_xy_ (Newson, 2002). It is also a normalized Mann-Whitney U statistic (Mason and Graham, 2002), which is used in the Mann-Whitney U test or Wilcoxon rank-sum test, a popular nonparametric alternative for a t-test. The latter connection is especially important because it means that by testing the statistical significance of a difference between two groups using the Mann-Whitney U test, one is also testing the statistical significance of AUC and vice versa (Figure 3A).

#### Quadratic error-based measures

Performance measures based on the quadratic error (together with information-based measures) quantify the performance of probabilistic predictions directly, without any intermediate transformation of predictions to categorical predictions or ranks. This makes them the most sensitive measures to capture signal in the data or model improvement, although it requires that the model predictions are in the form of probabilities.

Brier score (Brier, 1950) is the most prominent example from this category. It is a mean squared error between predicted values and observed values, where predictions are coded as 0 and 1, for the case of the binary classifier. The Brier score can be straightforwardly generalised to multi-class classification using ‘one-hot’ dummy coding.

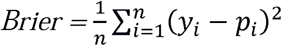

For example, if a model predicts that a patient with a disease has a disease with a probability of 0.8, the squared error will be (1-0.8)^2=0.04 and the Brier score is the average of error of all predictions across all subjects. Brier score has values between 0 and 1 (smaller the better), with 0.25 for a chance level predictions in a case where there is an equal number of subjects in both groups (because 0.5^2=0.25). Compared to AUC, it takes into account specifically predicted probabilities of an outcome, not only their ranks. The score is improved when the predictions are well calibrated, so when the predicted probabilities correspond to observed frequencies of misclassification (e.g. in subjects with the predicted probability of a disease of 0.8, 80% of these subjects will have the disease).

One of the difficulties with employing the Brier score in practice is that – unlike accuracy and AUC – it lacks a simple intuitive interpretation. In order to make it more intuitive, it can be rescaled to form a pseudo R^2^ measure analogous to variance explained used in evaluate regression models.

The Scaled Brier score is defined as

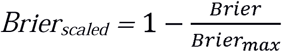

where the *Brier*_*max*_ is the maximum score that can be obtained with a non-informative model

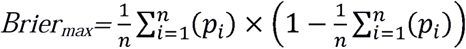

Thus a non-informative model will have a score of 0 and a perfect model score of 1, regardless of class frequencies.

One intuitively interpretable measure is a discrimination slope, also known as Tjur’s pseudo R^2^ (Tjur, 2009).

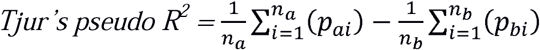

This is simply the difference between the mean of predicted probabilities of two classes, which can also be easily visualized (Figure 4).

**Figure 4:**
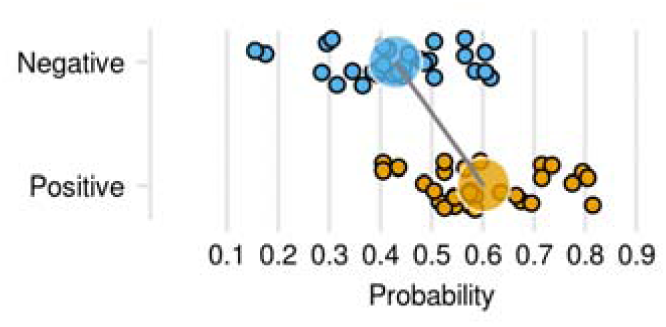
Tjur’s pseudo R^2^. Interpretation of Tjur’s pseudo R^2^ is the difference between mean predicted probability of the positive group and the negative group.

#### Information criteria

Information theory provides a natural framework to evaluate the quality of model predictions according to how much information about the outcome is contained in the probabilistic predictions. The most important information-theoretical measure is the logarithmic score, defined as

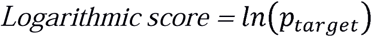

where p_target is the predicted probability of the observed target. If the target is coded as 0 and 1, and the score is averaged across all predictions, this becomes

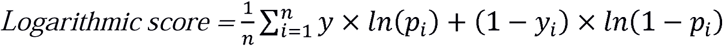

For example, a model predicts that a patient with a disease has a disease with a probability of 0.8, the logarithmic score will be ln(0.8) = −0.22. It has many interpretations and strong connections to other important mathematical concepts. It is a log-likelihood of observed outcomes given model predictions, it also equals to Kullback–Leibler or relative entropy between observed target values and predicted probabilities. It quantifies an average information loss (in bits) about the target given that we have only imperfect, probabilistic predictions. It also quantifies how surprising the observed targets are given the probabilistic predictions. In a certain sense, the logarithmic score is an optimal score. It has been shown that in betting or investing, the expected long term gains are directly proportional to the amount of information the gambler has about the outcome. For example, if two gamblers are betting against each other repeatedly on an outcome of football games, the gambler whose predictions are better according to the logarithmic score, will on average multiply her wealth, even if her predictions might be worse according to Brier score or AUC (Kelly, 1956; Roulston and Smith, 2000).

In practice, the logarithmic score and Brier score usually produce similar results and the difference is evident only with severe misclassification. The logarithmic score can grow to infinity when the predicted probability of an outcome is close to zero.

This might be considered undesirable since a single extreme wrong prediction can severely affect the summary score of an otherwise well-performing model. On the other hand, it can be argued that this is a desirable property of a score and predictions close to 0 and 1 should be discouraged. Intuitively, the logarithmic score measures surprise, if an event that is supposed to be absolutely impossible (p=0) happens, we rightly ought to be infinitely surprised. Similarly, if we know that an event is absolutely certain (p=1), then the right approach (mathematically) would be to bet all our money on this event, thus it is desirable that the wrong prediction is maximally penalized.

Similarly to the Brier score, the logarithmic score can be scaled to make it more intuitively interpretable. One popular way is to use Nagelkerke pseudo R^2^ (Nagelkerke, 1991) which is defined as

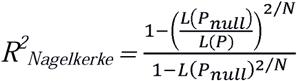

Where L(P) is a logarithmic score of model and L(P_null_) is a logarithmic score of chance level predictions (i.e. predicting only class frequencies).

### Empirical evaluation

Here we perform several empirical evaluations of statistical properties of the selected performance measures, one from each family described above, namely, accuracy, AUC, Brier score and logarithmic score. We examine: (i) the statistical power of detecting a statistically significant result, (ii) the power to detect a statistically significant improvement in model performance, (iii) the feature selection stability, and (iv) the reliability of cross-validation results. We employ real and simulated datasets.

#### Statistical power of detecting a statistically significant result

Statistical tests: In this section, we evaluate how often a given method is likely to detect a statistically significant effect (i.e., a classifier that exceeds chance levels at a given significance level, nominally p < 0.05). In this case, the focus is on statistical significance, but not on the absolute level of performance. We used the same datasets using the different performance measures described above. We tested the power of a model to make a statistically significant prediction on an independent test set. Statistical significance of the accuracy measure was obtained using a binomial test. To obtain statistical significance of AUC, we performed Mann-Whitney U test. Since AUC is equivalent to Wilcoxon-Mann-Whitney U statistics, performing Mann-Whitney-U test on model predictions equals to performing a statistical significance test of AUC. P-values for Brier score and log score were obtained using a permutation test, where the real labels were shuffled for a maximum of 10,000 times and the specific scores were computed for each shuffle, thus obtaining an empirical null distribution of scores where the specific p-value corresponds to a percentile of the observed test statistic in this distribution. For permutation tests, we employed early stopping criteria according to (Gandy, 2009), that stopped the permutation when the chance of making wrong decision to reject the null hypothesis was lower than 0.001. Performing the permutation test for accuracy and AUC is not necessary, because their distributions are known and can be computed exactly. These tests are only valid because we are testing the statistical significance in an independent test set. However, they would produce overly optimistic results in a cross-validation setting. This is because data points between folds are no longer completely independent and a permutation test where a model is refitted in each permutation should be used instead (Noirhomme et al., 2014; Varoquaux et al., 2017).

### Experiments

First, we examined statistical power on simulated distributions of model predictions, without any machine learning modeling. Similar to the first experiment, we repeatedly sampled from two one-dimensional Gaussian distributions 1 SD apart representing the distribution of model predictions of two classes. These predictions were transformed into 0-1 range using a logistic function and into categorical predictions by thresholding at 0.5 threshold. We performed 1000 simulations for sample sizes 20, 80, 140, 200, and recorded proportion of times each statistical test obtained a statistically significant result (p < 0.05).

Second, we examined statistical power of a support vector machine (SVM) to discriminate between two groups in the simulated dataset. The simulated dataset consisted of 6 independent variables, 3 signal variables and 3 noise variables. Signal variables were each randomly sampled from a Gaussian distribution with SD=1 and mean=0 for group 0 and mean=0.3 for group 1. Three noise variables were each randomly sampled from a Gaussian distribution with mean=0 and SD=1. We repeatedly sampled training and test set from this dataset of size 40, 80, 120, 140, and fit a support vector machine classifier in the training set and evaluated the statistical significance of the predictions in the test set. We used C-SVM implementation of an SVM with a linear kernel from a package kernlab (Karatzoglou et al., 2004), with a C parameter fixed at 1. SVM predictions were transformed into probabilities using Platt scaling (Platt, 1999), as implemented in kernlab.

Third, we examined statistical power on real neuroimaging datasets, including OASIS cross-sectional (Marcus et al., 2010) and ABIDE datasets (Craddock et al., 2009). We used already preprocessed OASIS VBM data as provided by the OASIS project using nilearn dataset fetching functionality (Abraham et al., 2014) The preprocessing consisted of brain extraction, segmentation of white matter and gray matter, and normalization to standard space using DARTEL. Details of the preprocessing can be found elsewhere (Marcus et al., 2010). Here we used gray matter and white matter data separately in order to predict biological sex and diagnostic status (presence or absence of dementia). To reduce the computational load, we reduced the dimensionality of the WM and GM datasets to 100 principal components each. Furthermore, we used already preprocessed ABIDE dataset as provided by the preprocessed-connectome-project (Craddock et al., 2009)to predict biological sex. These consisted of ROI average cortical thickness data obtained using ANTs pipeline (Das et al., 2009), defined using sulcus landmarks according to the Desikan-Killiany-Tourville (DKT) protocol (Klein and Tourville, 2012).

Additionally, we included common non-imaging machine learning benchmark datasets obtained from using mlbench and kernlab libraries originaly from UCI machine learning repository (Dua and Graff, 2017). These included Pima Indians diabetes, sonar, musk, and spam.

Finally, in order to compare statistical power to detect statistically significant above chance prediction according to specific performance measures, it is important to show that the higher power is due to higher sensitivity of the performance measure and not because the used statistical test is overly optimistic. To do this, we repeated all experiments including simulated and real datasets, but with permuting true labels before each simulation to destroy any relationship between the data and target outcome.

## Results

In all simulated and real datasets, when the null hypothesis was true (i.e. performing experiments on data with shuffled labels), statistical significance p < 0.05 was obtained approximately 5% of times for significance tests of AUC, Brier score and logarithmic score, as expected (supplementary figure 1). The binomial test was often overly conservative, i.e., p < 0.05 was obtained less than 5% of times. This is a known behavior of binomial test in small samples caused by a limited number of values in the null distribution (Fig 4 shows the example of n=20 sample). This conservativeness is worse in small samples and it disappears when the sample size is sufficiently large (i.e. N=5000). This behavior is not limited to a binomial test, it also happens if the p-value of accuracy is calculated using permutation test, which is just a random approximation of the exact binomial test, and the low resolution of the null distribution of accuracy results is present even when this distribution is obtained using permutations.

For all experiments and all sample sizes, tests using accuracy had the lowest power. Brier score, logarithmic score, and AUC performed approximately the same. At the sample size where AUC, Brier score, the logarithmic score obtained the common goal of 80% power, accuracy obtained only 60% power in all datasets (see figure 5). Results for all datasets separately are in supplementary figure 2

**Fig 5:**
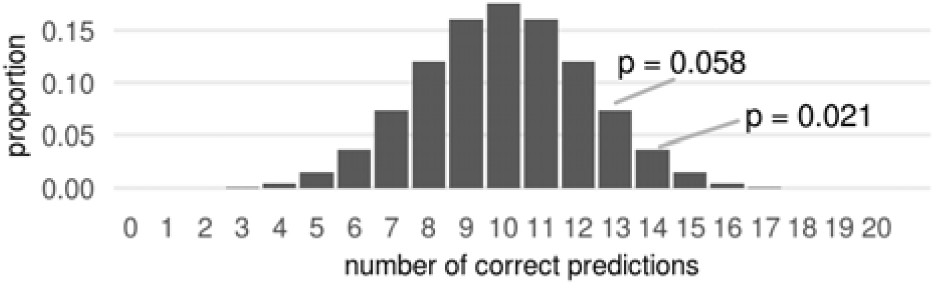
Null distribution of correct predictions for n=20 illustrating why significance test for categorical predictions is conservative in small samples at specific significance levels. Since the predictions are categorical, the null distributions consist only of a limited number of values. It is not possible to obtain p=0.05, in order to obtain p<0.05 it is necessary to get at least 14 correct predictions, corresponding to p=0.021. Although this makes the test conservative at the p = 0.05 level, it is not conservative at p = 0.058 or p=0.021 levels obtained by getting 13 or 14 correct predictions respectively.

### Power to detect a statistically significant improvement in model performance

In the previous section, we evaluated power to detect an effect that is significantly different from zero (i.e. exceeding ‘chance’ level), however, in many cases, it is important to detect a difference between two different classification models, potentially trained using different algorithms or different features. For example, we might want to know if a whole brain-machine learning model predicts remission of a depressive episode better than simple clinical data. Or we might want to compare the performance of two different methods e.g. support vector machines and deep learning. In both of these cases, it is important to evaluate the difference statistically, otherwise the apparent superiority of one model might be due to chance and translate to better predictions in a population.

#### Statistical tests

We obtained statistical significance of a difference in accuracy between two models using McNemar test using an exact binomial distribution. To test differences between two models using Brier score and logarithmic score, we used a permutation sign flip test testing the hypothesis that the difference in errors between two models is centered at 0. If the predictions were categorical, the sign-flip test would approximate the results of the exact McNemar test. There is no equivalent test for AUC because errors depend on ranks and thus cannot be computed for individual data points. Instead, we have used DeLong’s non-parametric test of differences between two AUC (DeLong et al., 1988).

#### Experiments

We compared the performance of two models on the same simulated and real datasets as in the previous section. Each time, we compared the performance of a model that was trained on the whole training set, with a model trained using only a subsample of the training set of size between 10-90% of the full training set. The model trained with fewer data points should eventually perform worse than a model using all available data. For each sample size of the restricted model, we have calculated the proportion of times a statistical test found a statistically significant difference in the performance of these two models.

#### Results

The difference between the power of different performance measures was higher than in the testing against the null hypothesis of no effect in the previous section. Accuracy had the lowest power, followed by AUC and Brier and logarithmic score performed approximately the same.

### Evaluation of stability of the feature selection process

Feature selection is an important part of machine learning with the goal of selecting a subset of features that are important for prediction. It is usually done in order to make models more interpretable and improve their performance. Different feature selection criteria lead to a different set of selected features. Here we evaluated which specific performance measure (accuracy, AUC, brier score, logarithmic score), leads to better feature selection results when its improvement is used as a criterion for feature selection. We performed a greedy forward stepwise feature selection, starting with 0 features and subsequently adding additional features into the model that improve the model performance the most according to a specific performance measure. This is a noisy feature selection process but preferably we would want informative features to be selected on average more often than non-informative features. We constructed stability paths according to Meinshausen Buhlman (Meinshausen and Bühlmann, 2010), the feature selection procedure was performed repeatedly on a random subsample of a dataset and the probability of selecting a specific feature was computed for any number of selected features in the final model. This was performed on 2 benchmark machine learning datasets 1, Pima Indians diabetes dataset and 2, spam prediction datasets. To each dataset, we added non-informative features by randomly permuting some of the original features, thus destroying any information they can have about the outcome. We compared how often original informative features are selected compared to non-informative features.

#### Results

Signal features (i.e. features that had not been permuted) were selected most often if the feature selection was performed according to logarithmic score and Brier score, followed by AUC and accuracy. For example for spam data, only one signal feature was stably selected using accuracy.

### Reliability of the cross-validation measure

Previously (Varoquaux et al., 2017) showed that cross-validation estimates of accuracy have high variance and high error with respect to the accuracy obtained on a large hold-out validation set, especially when the sample size is low. Here we compared the relationship between cross-validation performance and hold-out performance for accuracy, AUC, Brier score and logarithmic score. We performed a procedure similar to Varoquaux 2017. We selected 6 large sample datasets from UCI machine learning repository in order to have at least 1000 samples in the hold-out set. Further, we manipulated datasets by randomly flipping labels to 0-20% data points, thus creating many datasets with different true performance. In each of these datasets, we estimated model performance using 10-fold cross-validation and compared it to out of sample performance on the validation sample of size 1000. We repeated this procedure for different sizes of the training set 50, 100, 150, 200, and 250.

#### Results

In almost all comparisons, reliability of the cross-validation results (as measured by Spearman correlation between cross-validation performance and hold-out performance) was higher for logarithmic score and Brier score, compared to accuracy and AUC. This was true when the results for different datasets were combined together (Figure 7A) or separated (Figure 7B).

**Figure 6.**
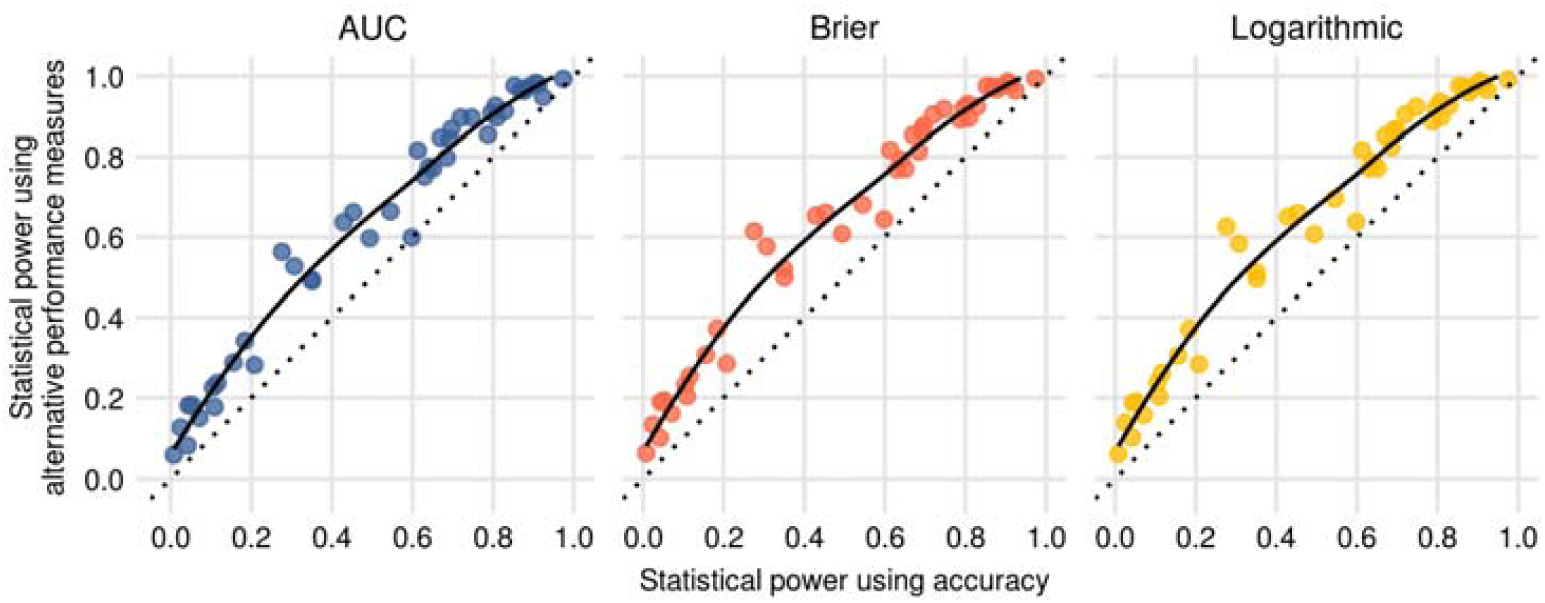
Comparison of statistical power of accuracy and alternative performance measures. We calculated statistical power to find significant above-chance performance in the test set (p < 0.05) across multiple simulated and real datasets with varying sample sizes. Each dot represents a proportion of statistically significant results across 1000 draws from a specific dataset and sample size. This figure shows that all alternative measures show greater power than accuracy for detecting a significant effect.

**Figure 7.**
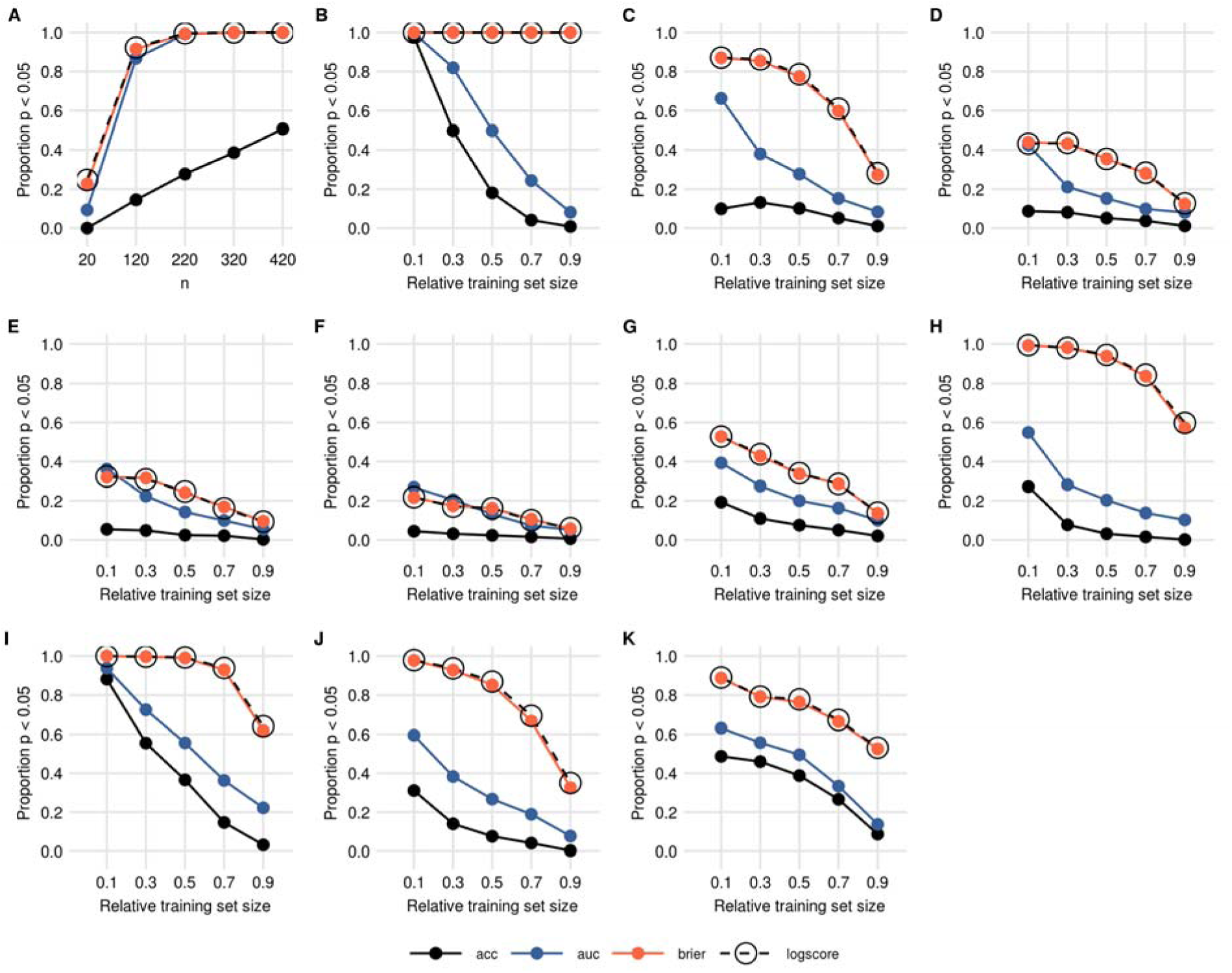
Statistical power of comparing 2 competing models. One model was trained using the whole training set and the second model was trained on fewer subjects, using only a proportion of the training set. Y-axis shows the proportion of statistically significant results obtain by comparing the performance of two models, from 1000 random draws from each dataset. A: simulated predictions, the performance of the two models was fixed, but we manipulated the sample size. B: SVM fitted on simulated data, C: OASIS gray matter gender prediction, OASIS white matter gender predictions, E: OASIS gray matter diagnosis prediction, F: OASIS white matter diagnosis prediction, G: ABIDE cortical thickness gender prediction, H: Pima Indians diabetes benchmark dataset, I: Sonar benchmark dataset, J: Musk benchmark dataset K: Spam benchmark dataset.

**Figure 8:**
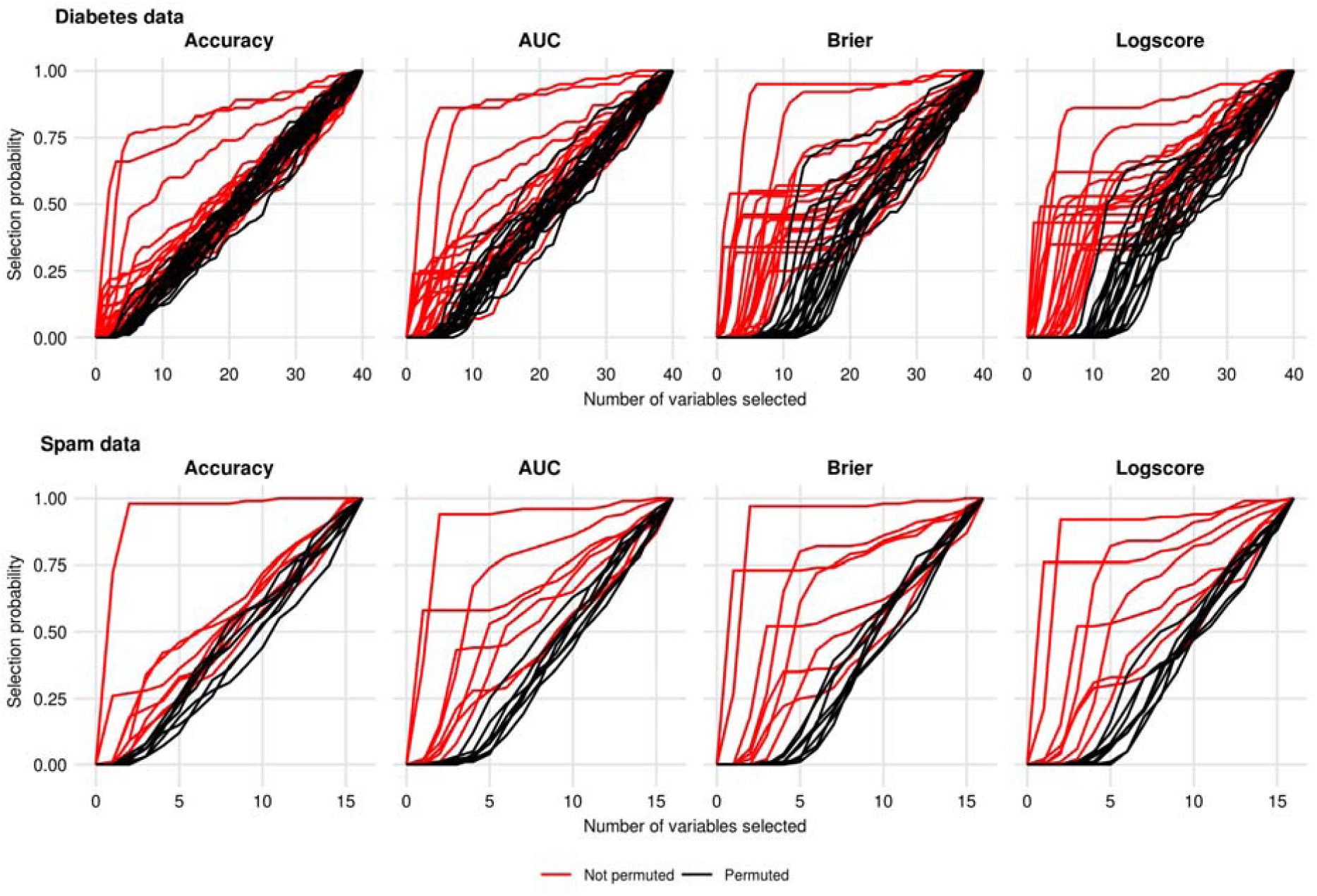
stability of feature selections. We can see stability paths of individual features if they are selected according to specific performance measures in a greedy forward stepwise feature selection procedure. Signal features (red) were selected sooner using logarithmic score and brier score compared to accuracy and AUC.

**Figure 7:**
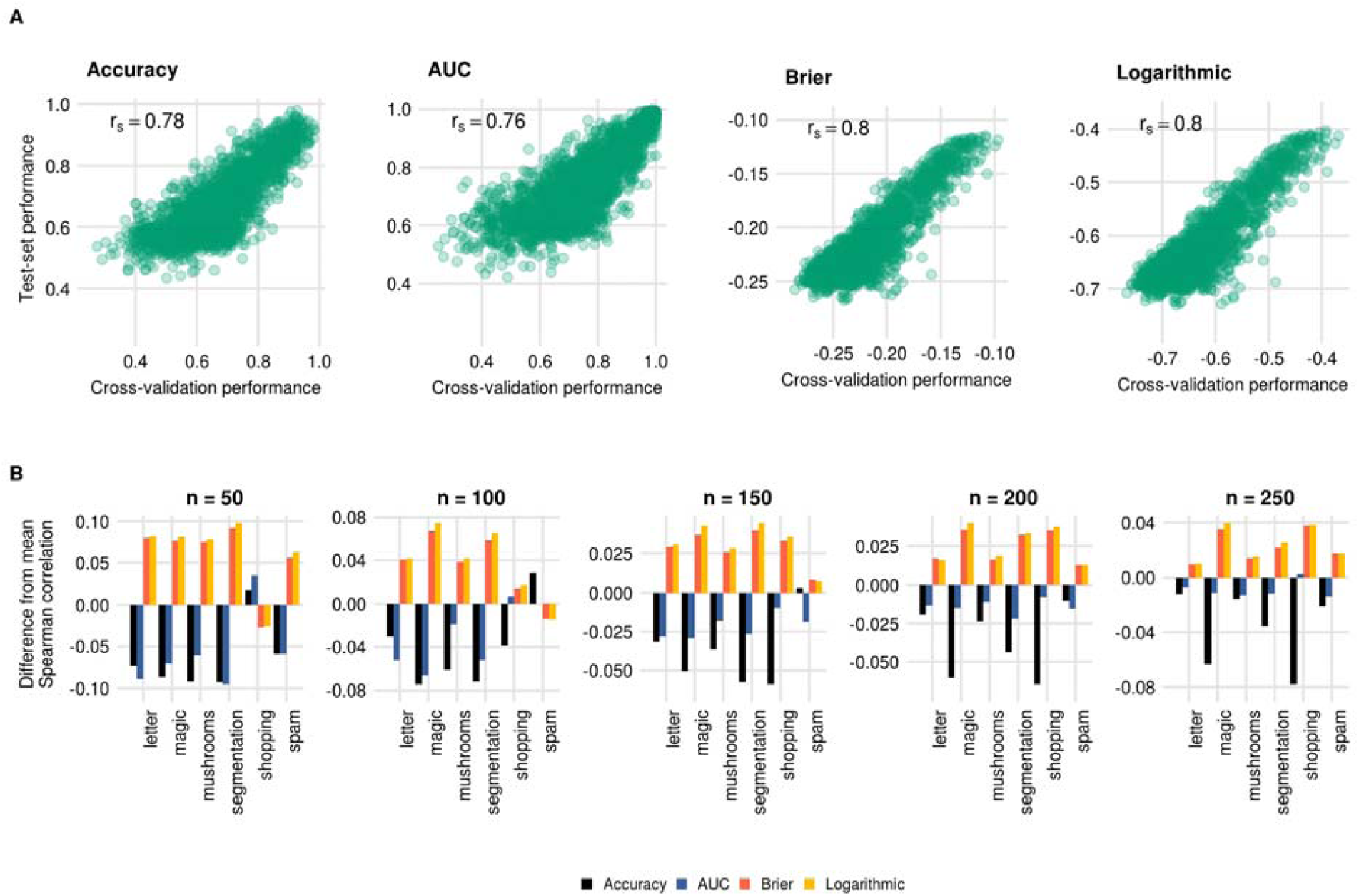
Comparison between performance in the cross-validation and on the holdout set. A: comparison across 6 datasets for a training set of size 50. B comparing Spearman correlation between cross-validation performance and hold-out performance for each sample size and dataset separately.

## Data and code availability

All data and code necessary to reproduce the experiments is at https://github.com/dinga92/beyond-acc

## Discussion and recommendations

The choice of a performance measure should be motivated by the goals of the specific prediction model. In this report, we reviewed exemplars of 4 families of performance measures evaluating different aspects of model predictions. Namely, categorical predictions, ranks of predictions and probabilistic predictions according to quadratic error and information content.

Often, such as in brain decoding and encoding, the primary goal of machine learning models is not to make predictions but to learn something about how information is represented in the brain. Two common goals of encoding or decoding studies are to establish if a region contains any information at all about the stimulus or behavior, and to compare the relative amount of information about stimulus between multiple ROIs (Naselaris et al., 2011).

In this case, the performance measure should be chosen to maximize the statistical power of a test and reliability of results. Accuracy, although the most commonly used performance measure, performed the worst in all statistical aspects we have examined in this study compared to alternative measures (i.e. AUC, Brier score, logarithmic score). It has the lowest statistical power to find significant results and to detect a model improvement, it leads to unstable feature selection and the results are least likely to replicate across samples from the same population. For these reasons, the accuracy should not be used to make a statistical inference, although if needed it can be reported as an additional descriptive statistic.

The loss of power when using accuracy is significant. Compared to alternative measures, the power to detect statistically significant results dropped from 80% to 60% or from 60% to 40% on average across simulated and real datasets. The reason for this is twofold. First, accuracy does not take the magnitude of an error into account, thus it is a very crude and insensitive measure of model performance. The model improvement can only be detected at the decision threshold, thus leading to severely suboptimal inference. Second, since accuracy evaluates only categorical predictions, it can only take a limited number of values, thus a null distribution is tabulated which leads to conservative p-values (Fig 4). The smaller the sample size, the worse this effect is.

The loss of power is even more prominent when the goal is to find statistically significant improvement on an already well-performing model. In this situation, both accuracy and AUC perform significantly worse than probabilistic measures (log score, Brier score). This effect is especially prominent when comparing already well performing models. When the discrimination between classes is already large, there is only a small chance that a model improvement would result in changing of ranks of predictions (for the change in AUC) or predictions crossing the decision threshold (for the change in accuracy). Even potentially important model improvements can be missed if the model is assessed based on accuracy. Situations are easily constructed where model performance improves significantly, but without ever changing the proportion of correctly classified samples. Or in other words, statistical power to detect an improvement in model performance according to accuracy can be effectively zero.

The crudeness of accuracy leads to another important problem, and that is a loss of reliability and reproducibility of the results, as shown in our comparison of cross-validation results. If we replicate the same analysis on a different sample from the same population, the results using accuracy will be less similar to each other than results using alternative measures. This, together with a significant loss of power, leads to less replicable results. It is important to note that there is no upside to these problems in the form of for example higher confidence in conservative statistically significant results using accuracy. As it was pointed out before (Button et al., 2013; Ioannidis, 2005; Loken and Gelman, 2017) low power necessary leads to overestimation of the found effect, low reproducibility of the results, and higher chance that the observed statistically significant effect size does not reflect the true effect size. (Varoquaux et al., 2017) pointed out that the error bars for accuracy under cross-validation are large, and that the results are much more variable than is usually appreciated. From our results, it is clear that a large portion of this variability is attributable to using accuracy as a performance measure, and that unfavorable statistical properties of cross-validation can be improved simply by using alternative performance measures.

In many situations, the statistical properties of a specific performance measure are not an important aspect of model evaluation. In a clinical context, the utility of model predictions for a patient or a clinician is far more important than statistical power. Model evaluation based on categorical predictions is in this situation inappropriate because categorical predictions hide potentially clinically important information from decision-makers, and they assume that the optimal decision threshold is known and identical regardless of patient or situation. In clinical settings, it is generally recommended (Moons et al., 2015) to evaluate models based on their discrimination, usually measured by AUC and calibration which measures how well the predicted probabilities matches observed frequencies. In situations where the optimal threshold is fixed and known or when the decisions need to be made fully automatically, without any additional human intervention, it is appropriate to evaluate model performance with respect to its categorical predictions. However, misclassification should be weighted according to relative the cost of false positive and false negative misclassification in order to properly evaluate the utility of the model. Accuracy weighs false positive and false negative misclassification equally (or according to class frequencies with balanced accuracy) which is almost never the case, thus it will lead to wrong decisions.

To present and visualize the model performance, multiple options exist. Good visualization should be intuitive and informative. Confusion matrices, although common, do not show all available information because they show only categorical predictions. This can be improved by plotting the whole distribution of predicted probabilities per target class in the form of histograms, raincloud plots, or dot plots as in figure 4. This directly shows how well the model separates target classes, it might reveal outliers or situations where the performance is driven by only a small subset of accurately classified data points. This can be accompanied by a calibration plot with predicted probabilities on the x-axis and observed frequencies on the y-axis with fitted regression curve, showing how reliable the predicted probabilities are. An additional commonly used visualization is a ROC curve. This is arguably less informative and less intuitive than plotting the predicted probability distributions directly, however, it can be used in specific situations where it is useful to visualize the range of sensitivities and specificities across different decision thresholds.

## Conclusion

We extensively compared classification performance metrics from four families using simulated and real data. In all statistical properties we evaluated, accuracy performed the worst, thus it should not be used as a metric to statistically evaluate model predictions. Summary measures based on probability predictions (i.e. logarithmic score or Brier score) performed the best and thus, we recommend the use of these measures instead of accuracy. For model interpretation and presentation, various measures can be reported at the same time, together with a graphical representation of model predictions. If the model is supposed to be used in practice such as in a clinical setting, summary measures are not enough. Rather, we recommend that the model be evaluated with respect to its discrimination power, calibration, and clinical utility. Accuracy should be avoided because it weighs false positive and false negative misclassification equally, or according to class frequencies but not according to consequences for a patient.

