## Supplemental figures for "Beyond accuracy: Measures for assessing machine learning models, pitfalls and guidelines"

**Supplementary information**

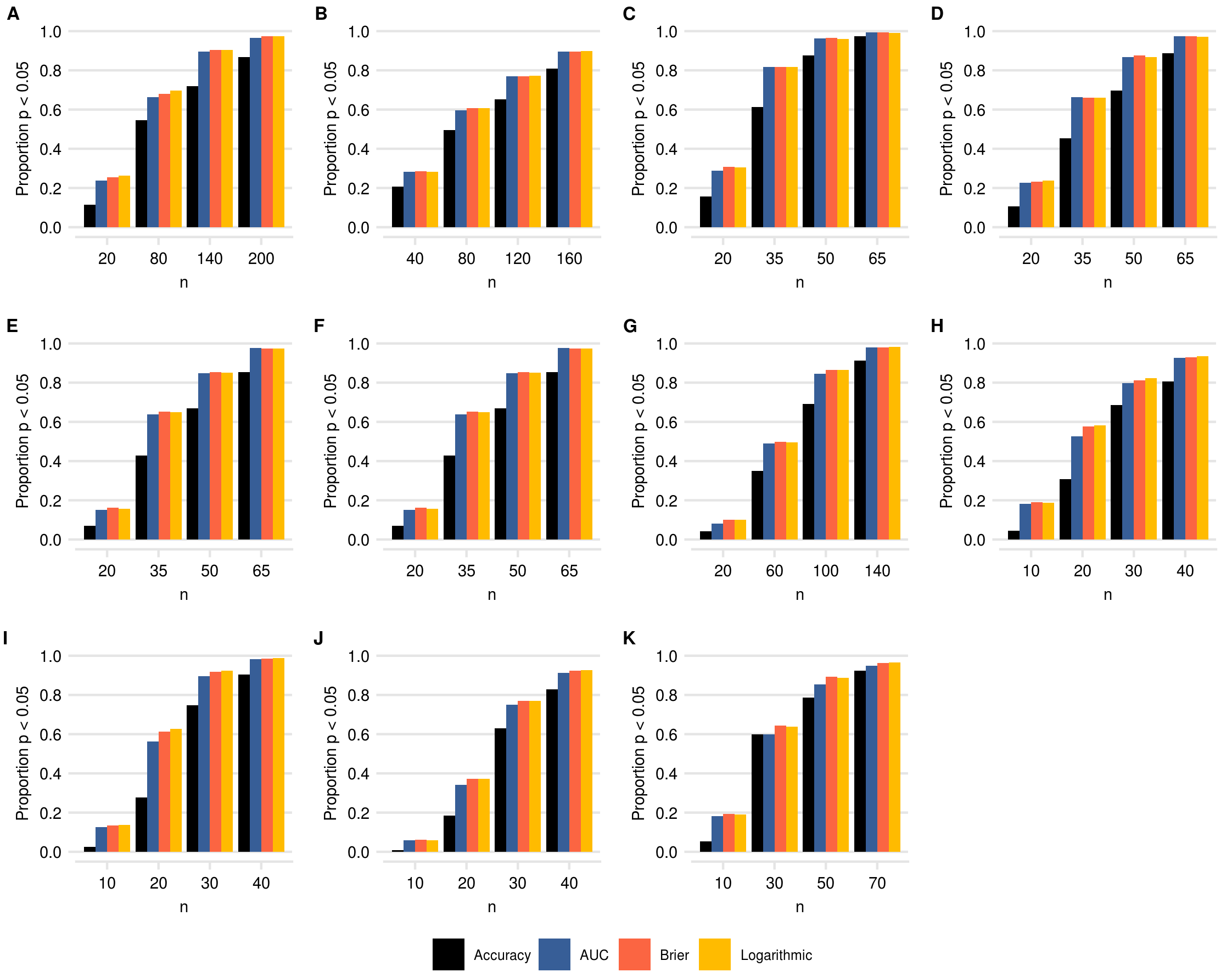

*Supplementary Figure 1: Statistical power of detecting above chance model performance. Y-axis shows the proportion of statistically significant results (p < 0.05) obtain from 1000 random draws from each dataset. A: simulated predictions, the performance of the the model was fixed, but we manipulated the sample size. B: SVM fitted on simulated data, C: OASIS gray matter gender prediction, OASIS white matter gender predictions, E: OASIS gray matter diagnosis prediction, F: OASIS white matter diagnosis prediction, G: ABIDE cortical thickness gender prediction, H: Pima Indians diabetes benchmark dataset, I: Sonar benchmark dataset, J: Musk benchmark dataset K: Spam benchmark dataset.*
